# Disentangling the importance of microbiological and physico-chemical properties of Ethiopian field soils for the Striga seed bank and sorghum infestations

**DOI:** 10.1101/2025.10.03.680328

**Authors:** Tamera Taylor, Getahun Benti, Marcio F.A. Leite, Desalegn W. Etalo, Lorenzo Lombard, Luisa Arias-Giraldo, Dominika Ryba, Stefan Sanow, Thomas Mostert, Einar Martinez de la Parte, José G. Maciá-Vicente, Daniel Yimer Legesse, Urguesa Tsega, Jiregna Daksa, Roy van Doorn, Raycenne R. Leite, Taye Tessema, Pedro W. Crous, Dorota Kawa, Eiko E. Kuramae, Jos M. Raaijmakers, Siobhan M. Brady

**Affiliations:** Department of Plant Biology and Genome Center, University of California, Davis, Davis, CA 95616; Department of Microbial Ecology, Netherlands Institute of Ecology (NIOO -KNAW), Droevendaalsesteeg 10, 6708 PB Wageningen, The Netherlands; Westerdijk Fungal Biodiversity Institute (WI-KNAW), Uppsalalaan 8, 3584 CT Utrecht, The Netherlands; Ethiopian Institute of Agricultural Research, Holeta, Ethiopia; Plant Hormone Biology Group, Swammerdam Institute of Life Science, University of Amsterdam, Amsterdam, Netherlands; Wageningen University, Laboratory of Phytopathology, Wageningen, The Netherlands; Marine Sciences and Applied Biology, University of Alicante, Alicante, Spain; Addis Ababa University, Addis Ababa, Ethiopia; Experimental and Computational Plant Development, Utrecht University, Padualaan 8, 10 3584 CH, The Netherlands; Plant Stress Resilience, Utrecht University, Padualaan 8, 10 3584 CH, The Netherlands; Ecology and Biodiversity, Institute of Environmental Biology, Utrecht University, Padualaan 8, 10 3584 CH, Utrecht, The Netherlands; Institute of Biology, Leiden University, 2333 BE Leiden, The Netherlands; Howard Hughes Medical Institute, University of California, Davis, Davis, CA 95616

## Abstract

*Striga hermonthica* (Striga) is a parasitic weed that severely affects sorghum yields in sub-Saharan Africa. Recent studies highlighted the soil microbiome’s potential to suppress Striga through interference with specific stages in its life cycle. In this study, meta-analysis of 48 Ethiopian field soils revealed that microbial communities and their interactions with soil physico-chemical properties correlated with Striga field occurrence. Striga infestation of sorghum and soil seedbank levels were negatively correlated with clay content and the nutrients potassium, sulfur, calcium, and carbon. Microbiome analyses indicated that fungal communities were more responsive than bacteria to changes in Striga infestation and seedbank levels, with distinct microbial compositions even in soils where Striga was not detected. Specific fungal and bacterial genera showed both positive and negative correlations with Striga measures, but patterns rarely held across taxonomic levels, highlighting the complexity of microbiome–Striga interactions. To begin to validate these correlations, we tested an isolate from the fungal genus *Neocosmospora*, which negatively correlated with the Striga seedbank, and showed that this isolate promotes Striga seed germination *in vitro*, suggesting potential for biological control of Striga. The data and analysis methods are integrated and shared in a public Shiny App for broader analysis and continued research on soil-Striga interactions.

## Introduction

*Striga hermonthica* (Striga), or witchweed, is a devastating agricultural weed for cereal and legume crops throughout the world [1,2]. For resource-poor farming communities in sub-Saharan Africa, Striga causes substantial yield losses of the staple crop *Sorghum bicolor*. Sorghum exudes strigolactones to recruit arbuscular mycorrhizal fungi (AMF), but this signal molecule is hijacked by Striga for seed germination [3,4]. Upon perception of other host-derived exudates, the Striga radicle forms a haustorium which penetrates the host root. Parasitism is complete once the Striga forms a xylem-xylem connection, through which Striga extracts essential nutrients and water from the host. Once Striga grows above ground, it is able to photosynthesize to complete its life cycle through the production of tiny seeds which are easily dispersed in the top soil layer [5,6]. Striga’s high fecundity, exceptional seed dormancy, easy seed dispersal, tight linkage to its host physiology, and a lack of consistently effective management strategies poses an incredible agricultural challenge in Striga endemic areas.

Several approaches have been developed that can partially suppress Striga parasitism of sorghum. These include resistance breeding [1,7,8], chemical approaches [9,10] and farming practices [11,12]. These approaches are not singularly effective, not necessarily financially accessible, nor is resistance breeding easily applicable to local, preferred crop varieties. Hence, an integrated management strategy is needed. In the past decade, interest in the functional potential of the soil microbiome emerged as a novel and complementary resource for parasitic weed management. The suppressive activity of the soil microbiome was exemplified in the recent study by Kawa et al. (2024) where the microbiome complement of a soil suppressed Striga infection of *Sorghum bicolor*, through modulation of the production or breakdown of haustorium-inducing factors [13]. Other studies focusing on single microbial taxa, such as the fungus *Fusarium oxysporum*, have shown reductions of Striga germination, emergence, and infection rates [14–16]. Collectively, these and other efforts demonstrate the largely unexplored potential of microbes to disrupt the parasite’s life cycle and its interaction with the host plant.

To further disentangle the importance of soil chemistry and the microbiome in Striga suppression, we profiled 48 Ethiopian field soils from 14 diverse agro-ecological zones for Striga infestation levels of sorghum, Striga seed bank, soil physico-chemical properties and microbiome composition. We identified the fungal soil community as the most sensitive to variations in Striga seedbank and Striga infestation levels monitored in the field. Also, interactions between physico-chemical properties, bacterial and fungal abundances were shown to collectively inform Striga prevalence. These data were compiled in a publicly available Shiny App that facilitates multifactorial analyses of fungal, bacterial and physico-chemical properties across the sorghum belt in Ethiopia.

## Results

### Striga occurrence and soil physico-chemical properties across Ethiopia

Forty eight soils were sampled across the sorghum belt in Ethiopia to maximize diversity in Striga presence, physico-chemical properties, and microbial community composition (**Figure 1)** [17]. Striga infestation (plants per m^2^, Striga/m^2^) and seedbank (Striga seeds per 150 g of soil, seeds/150g soil) varied considerably across the soils and locations, with soils containing no Striga infestation (samples E13, E16, E20, E21, E33, E34, E42, and E47) or seedbank (samples E13, E16, E17, E19, E20, E21, E30, E33, E38, E40, E43, and E45) while others presented with maximum infestation of 185.9 Striga/m^2^ (sample E12) and 85.5 seeds/150g soil seedbank (sample E22) (**Figure 2A-B, S1, S2**). The Kewet region is notable for its differences in agro-climatic zones and Striga occurrence despite close geographical proximity. The thirteen sites in this region were within 80 km of each other and represent four different agro-climatic zones (**Figure 1B**). Within the Kewet region, in agro-climatic zone 12, soils E03, E07, E08, E30 and E43 have a highly variable Striga seedbank and infestation (25.1, 3.4, 11.9, 0, 0 average seeds/150g soil and 118, 1.6, 18.4, 0, 10.1 average Striga/m^2^, respectively) (**Figure S1**). In contrast, soils E01, E02 and E19, spanning an 8 km distance and within the same agro-climatic zone (zone 4), have relatively small seedbank and infestation levels (0.9, 1.9, 0 average seeds/150g soil, and 0, 0, 1.8 average Striga/m^2^) (**Figure S1**). Variation in Striga infestation relative to the seedbank within similar geographic locations is also present. For example, soils E24 and E37, also in agro-climatic zone 4, are spaced apart by only 7 km and while they have similar seedbank levels (averaging 2.6 and seeds/150g soil, respectively), they differ in infestation values (averaging 68.5 and 28.5 Striga/m^2^) (**Figure S1**).

**Figure 1.**
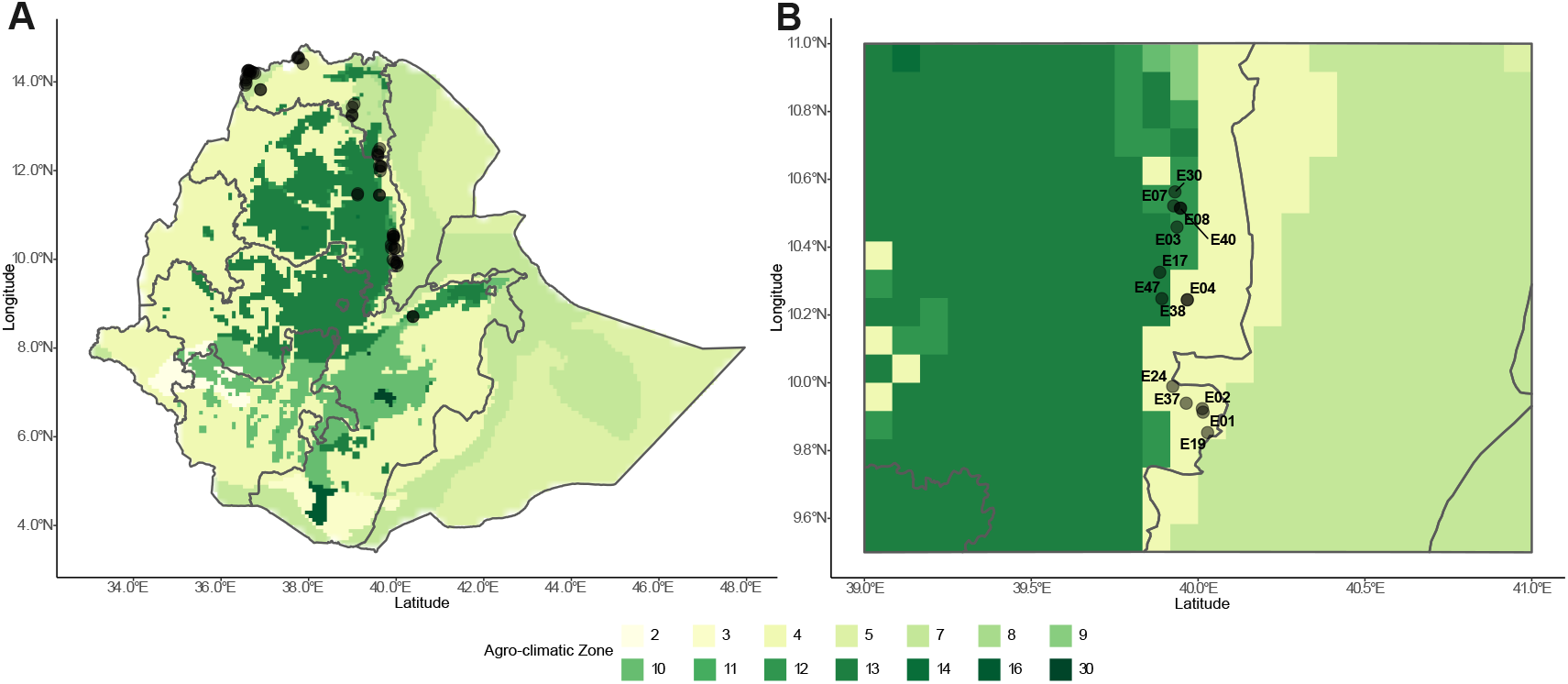
Ethiopian soil samples represent a diversity of agro-climatic zones. **A**. Map indicating locations of soil samples (black dots) as well as Ethiopian agro-climatic zone (green shades in heatmap) based on the 3 character Koeppen-Geiger climate classification using climatic data from CLUST32 (years 1981-2010) (source: GAEZ v4) (Table S1). **B**. Magnification of the Kewet region demonstrating the distance between samples in areas with varying soil and Striga infestation and seedbank. Koeppen-Geiger classifications 2- Equatorial monsoon, 3- Equatorial savannah, dry summer, 4- Equatorial savannah, dry winter, 5- Desert climate, hot, 7- Steppe climate, hot, 8- Steppe climate, cold, 9-Temperate/mesothermal climate, fully humid, hot, 10-Temperate/mesothermal climate, fully humid, warm, 11-Temperate/mesothermal climate, fully humid, cold, 12-Temperate/mesothermal climate, dry summer, hot, 13-Temperate/mesothermal climate, dry summer, warm, 14-Temperate/mesothermal climate, dry summer, cold, 16-Temperate/mesothermal climate, dry winter, warm, 30-Tundra climate.

Six distinct soil groups were defined based on hierarchical clustering of physical and chemical values (**Figure 2A**). Chemical properties drove differences between certain clusters. Cluster 1 is characterized by collective low levels of calcium (average 35.3%) and features samples with distinct chemical properties. E13 contains a high potassium content (7.4%, 18.3 mmol/kg) while E36 had a very high percentage of total magnesium (64%, 156.8 mmol/kg). E43 had a high percentage of potassium (6.4%) and magnesium (59%), as well as a high carbon to sulfur ratio (375). Cluster 3 shows an abundance of carbon (average organic and inorganic 2.9%) and a high available calcium (average 4.5 mmol Ca/L). Physical characteristics also inform the clustering with Cluster 3 being carbonated lime-heavy (average 4.9%) and Cluster 2 containing samples that have low clay, clay humus, and organic matter (average 24%, 180 mmol+/kg, 2.1%). Physical proximity in some cases was associated with similar composition. For instance, the physically proximal E01, 02 and 19 had similar levels of calcium, sulfur, inorganic carbon, organic matter and carbonated lime and clustered together in Cluster 3 (**Supp. Data 1 Table 1**). Other samples in Kewet; E04, E08 and E40 clustered separately into Cluster 6 and have similar levels of phosphorus, sulfur, potassium and nitrogen. In contrast, the geographically distant sample E14 also clusters in Cluster 6 due to its similar physico-chemical composition to E04, E08 and E40. The largest cluster, Cluster 4, had median levels of physico-chemical properties. Principal Component Analysis (PCA) was used to identify variables that explained the most variation in soil physico-chemical profiles. Together, PC1 and PC2 explained 41.5% of the variation (**Figure S3A**). The most influential factors in the physico-chemical PCA were total sodium, available potassium, and available sodium for PC1 and total sulfur, available calcium and total calcium for PC2 (**Supp. Data 1**).

**Figure 2.**
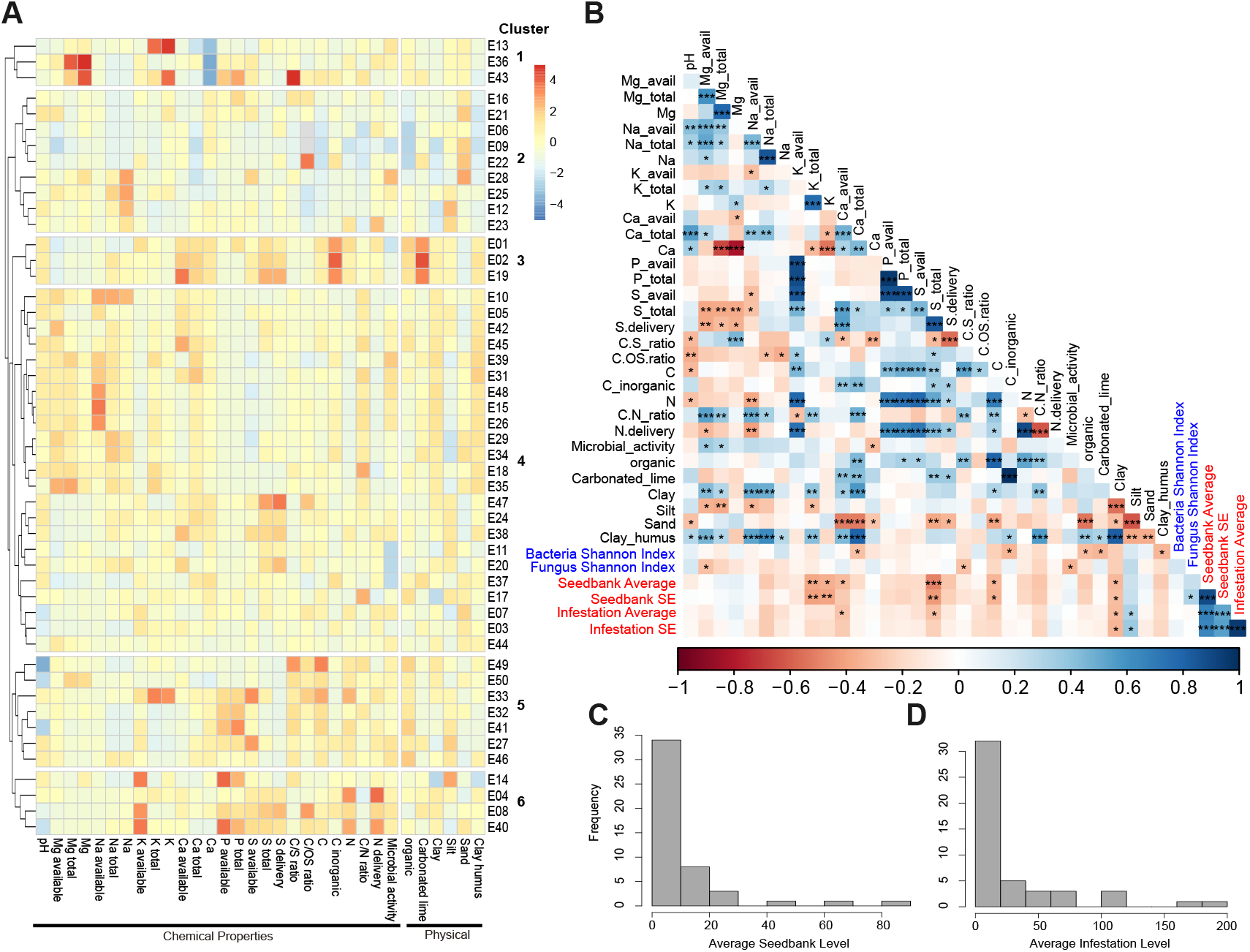
Soil samples vary in Striga infestation and seedbank levels, physicochemical parameters and microbial diversity. **A**. Heat map of soil physico-chemical properties and Striga measurements of all naturally infested soils excluding outlier sample 30. Color represents the relative value across samples (pheatmap). Soil samples were sorted by hierarchical clustering using a Euclidean distance metric with a tree cut off at 6 branches. **B**. Pairwise complete correlation analysis of physico-chemical soil factors, microbe diversity, and Striga occurence measurements. Color indicates correlation value as calculated by “psych” package in R. Significance levels 0.05, 0.01, and 0.001 represented by *, **, and *** respectively. Striga parameter labels highlighted in red. **C**. Distribution of Striga seedbank as seeds per 150 g of soil in soil samples. **D**. Distribution of Striga infestation as the number of Striga in a square meter normalized by the number of sorghum plants.

### Soil properties associated with Striga infestation and Striga seedbank

We next sought to determine if any soil physico-chemical parameters were significantly correlated with Striga infestation. Notable physical characteristics are clay and silt, which correlated with Striga measures in opposite manners. Clay negatively correlated with all Striga measures (seedbank r = −0.323, p = 0.024; infestation r = −0.31, p = 0.03), while silt showed positive correlations (seedbank r = −0.21, p = 0.15; infestation r = −0.31, p = 0.03). From the chemical analysis, relationships with Striga were found for potassium (K), calcium (Ca), sulfur (S), and carbon (C). Negatively correlating with Striga seedbank amount are total K (r = −0.38, p = 0.007), percent of K in overall nutrient content (r = −0.36, p = 0.013), total S (r = −0.49, p = 0.0004), and percent C in nutrient content (r = −0.34, p = 0.019). Total S also negatively correlates with Striga infestation (r = −0.29, p = 0.04). Available Ca negatively correlates with Striga seedbank (r = −0.31, p = 0.03) and infestation (r = −0.29, p = 0.048) (**Figure 2B**). Striga seedbank positively correlated with Striga infestation (r = 0.67, p = 0.0001) affirming the value of using seedbank as a proxy for infestation [17].

### Fungal communities change with Striga occurrence

The microbiome of these soils was profiled by 16S rRNA and ITS amplicon sequencing, capturing 19,905 and 10,749 unique bacterial and fungal amplicon sequence variants (ASVs), respectively (**Supp. Data 2)** [18]. Soils had variable bacterial (**Figure S2B**) and fungal (**Figure S2C**) phyla abundances, dominated by the bacterial phyla Acidobacteria, Actinobacteria and Proteobacteria and the fungal phyla Ascomycota, Basidiomycota, and unidentified phyla. Alpha diversity of bacterial and fungal communities was estimated with the Shannon Index, showing little variability in the bacterial communities across soils (6.5 ± 0.2), and more variability in the fungal communities (4.6 ± 0.4). (**Figure S1E-H, Supp. Data 2**). While the overall level of microbial diversity did not vary dramatically across samples, their community composition may significantly associate with Striga occurrence. The results showed that bacterial community diversity did not display a significant relationship with either Striga infestation or Striga seedbank values, however, fungal community diversity positively correlated with Striga seedbank variance (r = 0.30, p = 0.04) (**Figure 2B, Supp. Data 3**). PCA analysis of the ASV data for each microbial kingdom revealed no specific taxa as the most informative (**Supp. Data 2**). Bacterial community variation was only explained at 8.9% by PC1 and PC2 (**Figure S3B, Supp. Data 2**). The top ASVs driving PC1 are from multiple phyla with some redundancies at lower taxonomic levels but no clear lineage driving community composition (**Supp. Data 2**). For fungal ASVs, the top 2 PCs only explained 8.2% of community variation (**Figure S3C, Supp. Data 2**). Of the top 10 ASVs for fungal PC1, 8 were from unidentified phyla (**Supp. Data 2**). This suggests we may be capturing novel or taxonomically poorly resolved taxa contributing strongly to variations in the Striga seedbank and/or infestation.

Generalized Joint Attribute Modeling (GJAM) [19,20] revealed sensitivity of microbial groups microbes (bacteria+archeae and fungal communities) to changes in Striga. Striga infestation and seedbank variables were grouped into four discrete categories to represent zero, low, medium and high levels (**Figure 3**). The three different groups (bacteria/archae, fungi, physico-chemical variables) were submitted to sensitivity analysis with respect to changes in Striga infestation (**Figure 3A**) and seedbank (**Figure 3B, Supp. Data 3**). The fungal community displayed the highest sensitivity values, followed by the bacterial/archae community and soil physico-chemical factors. Interestingly, the sensitivity of fungal and bacterial groups (or communities) increases when the Striga infestation surpasses 40 Striga/m^2^ and the Striga seedbank surpasses 10-20 seeds/150g soil. However, for the Striga seedbank, we noticed that the microbial communities (fungal and bacterial) and soil physico-chemical factors still demonstrated relatively high sensitivity values for the sites with 0 Striga seedbank (**Figure 3B**). This suggests that the soils without a Striga seedbank still harbor a distinct microbiome.

**Figure 3.**
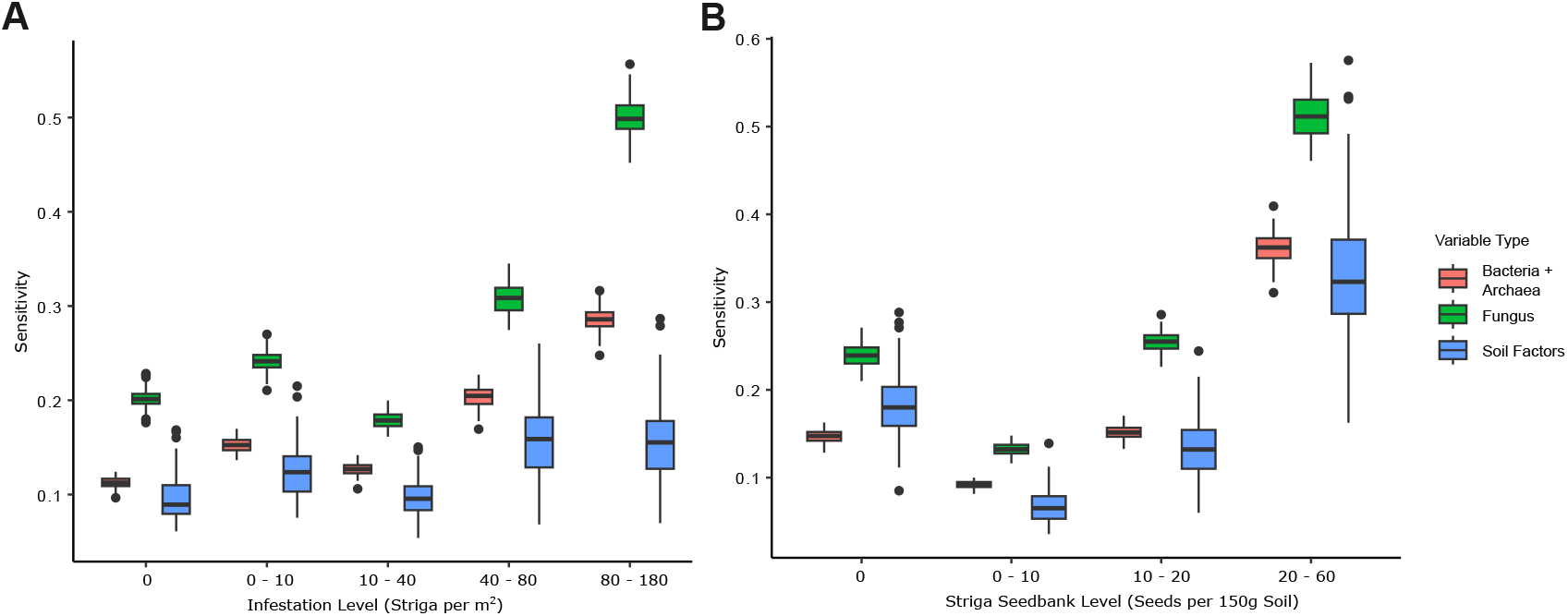
Soil fungal community composition changes the most with varying Striga occurrence. **A**. Sensitivity analysis of Striga field infestation against variable categories (bacteria and archaea, fungus, and soil physico- chemical factors). **B**. Sensitivity analysis of the same categories against Striga seedbank levels.

### Specific taxa correlate with Striga Infestation and the Seedbank

We next determined how specific fungal or bacterial taxonomic groups, from phylum through genus, related to Striga infestation, seedbank levels, or their variability by Pearson correlation (**Figure 4A-B, Supp. Data 4**). The analyses revealed that correlations with Striga shared between nested taxonomic levels were rare. For instance, despite the phylum-level negative correlation of Myxococcota with Striga field infestation (r = −0.32, p = 0.03), when looking at bacterial composition at the genus level, only one of 17 identified genera demonstrated a significant correlation with Striga (**Supp. Data 4**). *Cystobacter* was the only genus of Myxococcota with significance, yet it had positive correlations for seedbank level (r = 0.40, p = 0.006), demonstrating how much variation can exist within a single taxonomic group (**Figure 4A, Supp. Data 4**). Bacterial genera: *Lacunisphaera, Alterococcus* and *Cephaloticoccus* of the phylum Verrucomicrobiota, demonstrated a consistent negative relationship with Striga within a phylum (**Figure 4A**). *Alterococcus* negatively correlated with seedbank level (r = −0.30, p = 0.04), *Cephaloticoccus* negatively correlated with seedbank variance (r = −0.30, p = 0.04) and *Lacunisphaera* negatively correlated with Striga infestation (r = −0.29, p = 0.05) (**Supp. Data 4**). While these genera were all of the same family, Opitutaceae, the variable abundance of other genera and the unknown ASVs in this taxon was such that an association with Striga was lost. In fact, none of the higher taxonomic groups of this clade; order Opitutales, class Verrucomicrobiae, or phylum Verrucomicrobiota, showed a correlation with Striga (**Supp. Data 4**). With the amount of variation in microbiological life strategies, different associations within a taxonomic level are to be expected.

**Figure 4.**
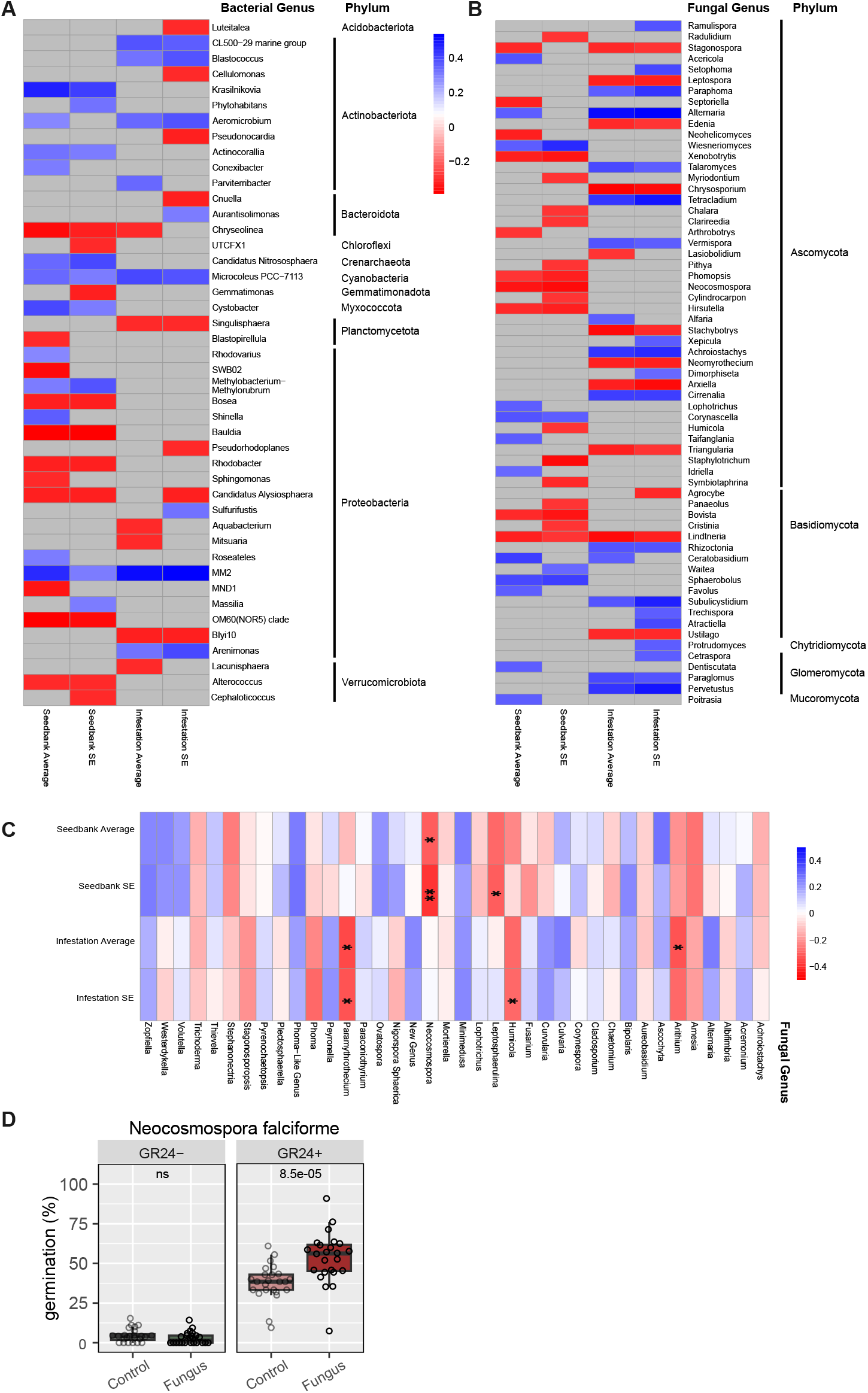
Specific microbial genera negatively correlate with Striga occurrence in soils. **A**. Bacterial genera with significant correlations to Striga occurrence (Pearson, non-adjusted p-value <0.05). **B**. Fungal genera with significant correlations to Striga occurrence (Pearson, non-adjusted p-value <0.05). Genera of both plots are grouped by phylum for visualization purposes. Color scale is shared for both plots A and B. **C**. Pearson correlation results of all reassigned fungal genera with Striga occurrence, displayed in alphabetical order. Significance levels of non-adjusted p values 0.05, 0.01, and 0.001 represented by *, **, and *** respectively. **D**. A strain of the species Neocosmospora falciforme tested for effect on Striga seed germination in vitro with and without the addition of synthetic strigolactone GR24. Statistics presented represent p-values from a wilcoxon test comparing means within GR24 treatment.

Both positive and negative relationships were identified between fungal genera and Striga. Genera of the Glomeromycota phylum with correlations to Striga were each in the positive direction (**Figure 4B**). These genera represent two different classes within this phylum, Glomeromycetes and Paraglomeromycetes. In the former, *Cetraspora* correlated with infestation variance (r = 0.30, p = 0.04) while *Dentiscutata* correlated with seedbank (r = 0.31, p = 0.038). Of the latter, *Paraglomus* and *Pervetustus* both correlated positively with infestation levels (r = 0.34, p = 0.022; r = 0.38, p = 0.009) and its variance (r = 0.33, p = 0.026; r = 0.43, p = 0.002) However, neither of these classes correlated with Striga on their own (**Supp. Data 4**). Within the Agaricomycetes class of Basidiomycota are many genera associated with Striga occurrence, although in opposing ways, resulting in neither significantly correlating with Striga. Genera with negative correlations to Striga seedbank were in the Agaricales order; *Agrocybe* (infestation SE r = −0.31, p = 0.039), *Panaeolus* (seedbank SE r = −0.29, p = 0.049), *Bovista* (seedbank r = −0.31, p = 0.036), *Cristinia* (seedbank SE r = −0.29, p = 0.047), and *Lindtneria* (seedbank r = −0.33, p = 0.027; infestation r = −0.3553, p = 0.0154). Even so, the Agaricales order itself did not show a significant correlation with Striga due to other ASVs in this order with abundance differing from this trend (**Supp. Data 4**). However, order Cantharellales in the Agaricomycetes class positively correlated with seedbank levels (r = 0.32, p = 0.029) and is bolstered by three of seven identified genera in Cantharellales, also demonstrating positive correlation with Striga; *Rhizoctonia* (infestation r = 0.32, p = 0.030), *Ceratobasidium* (seedbank r = 0.35, p = 0.016; infestation r = 0.30, p = 0.044), and *Waitea* (infestation r = 0.27, p = 0.07). Other genera of the Agariomycetes class also showed positive correlations with various Striga measures. *Subulicystidium* and *Trechispora* had positive correlations with Striga infestation (r = 0.40, p = 0.006) which coincides with their order, *Trechisporales*, correlating as well (r = 0.43, p = 0.003). Taxonomic groups that demonstrate a consistent negative relationship with Striga at multiple levels represent groups of species that may prove fruitful to investigate further for diminishing the Striga seedbank or reducing infection.

### Can individuals from potentially suppressive microbial genera directly impact Striga seed

As a proof-of-concept, we aimed to test the capacity of a representative isolate from a microbial genus which was negatively correlated with Striga occurrence. To better resolve their taxonomy, ASVs were re-assigned to the genera of individually cultured Ethiopian soil fungal isolates who had the highest bit score match [21] (**Supp. Data 5**). One of these re-assigned genera was *Neocosmospora* which showed a significantly negative correlation with Striga seedbank (r = 0.32, p = 0.03). A strain of *Neocosmospora falciforme* was tested *in vitro* for its impact on Striga seed viability and germination and demonstrated increased Striga seed germination (**Figure 4D, Supp. Data 5**). Promotion of germination under unsuitable conditions in absence of the host, also referred to as suicidal germination, is a highly promising strategy to diminish the Striga seedbank over time [22].Similarities between the two studies demonstrate a possibility of certain taxa, like *Neocosmospora*, perturbing the Striga seedbank or host infection in multiple experimental conditions. It should be noted, however, that *in vitro* settings may not always recapitulate the microbial dynamics and activities in complex conditions that exist in the field and therefore it is likely to also find mismatches between the results of our computational and experimental approaches.

### Provision of data for research and agricultural communities - Shiny app

Analysis strategies from this study, as well as other approaches were integrated into an interactive application (https://brady-lab.shinyapps.io/EthiopianSoil/). The purpose of this ShinyApp is to make the data freely accessible and easily analyzed for those who work with Sorghum breeding, pest management, microbiologists, and soil scientists within Ethiopia, sub-Saharan Africa, and beyond. This app is equipped with all foundational data described here, including soil physico-chemical and Striga measurements, and microbiome ASV counts and taxonomy (**Supp. Data 1-3**). As an example, a user can explore, in a visually interactive manner, soil physico-chemical and microbial phyla associated with the highest Striga field infestation. In the “Map View” window, one can select Striga infestation (Normalized_Striga_Count_Ave) to be plotted. The color scale indicates the degree of Striga infestation, with soil E12 having the highest Striga field infestation of 185.9 Striga/m^2^ (**Figure 5A, S2, Supp. Data 1**). In the “Individual Soil Compositions” window, selection of E12 provides an overview of soil physical and nutrient parameters as well as microbial composition, broken apart by bacterial and fungal phyla (**Figure 5B**). All graphs are easily downloadable. One can also perform analyses of multiple samples to determine correlations (using either Pearson or Spearman correlation as a distance metric) with any of the physicochemical, microbial and Striga infection parameters. Data can be summarized in a heatmap, used to demonstrate relationships through Pearson and Spearman correlations, or lines of best fit, and even simplified in a PCA. These strategies allow users to focus on traits or samples most interesting to them based on their own work or prior knowledge. They can have background information for excluding samples from certain areas or only want to look at relationships between certain chemicals. The methods included here can demonstrate patterns or abnormalities to consider in further analysis. For instance, using the app to create a PCA plot for all samples based on chemical analysis data reveals how sample E30 does not cluster regularly with the rest of the soils (**Figure 5C**). Observation with the heatmap confirms that E30 is not representative of the other samples due to a six-fold increase from the average in values for phosphorus, potassium, and sulfur measured (**Figure 5D**). This sample can easily be removed from the analysis by deselecting it from the sample pull down menu. Microbe data is included in a separate selectable section to walk through data preparation at different taxonomic levels before combining with other user-selected soil data. Correlation analysis can be run at this level to reveal relationships not explored in this text. For instance, in looking at fungal phyla correlation with soil chemical measurements, we see phylum Glomeromycota (representative arbuscular mycorrhizal fungi, AMF), in which many genera associated positively with Striga (**Figure 4B, Supp. Data 4**), also shows a negative relationship with available and total phosphorus (r = −0.39, p = 0.007; r = −0.38, p = 0.008) (**Figure 5E**). With these capabilities, users can explore the dataset and download data, analysis and figures, depending on their own research questions and insights. Future efforts will be geared towards improving soil-Striga modeling with additional data so that it may be used for predicting Striga pressure and mitigation techniques based on user inputted information such as location or soil parameters.

**Figure 5.**
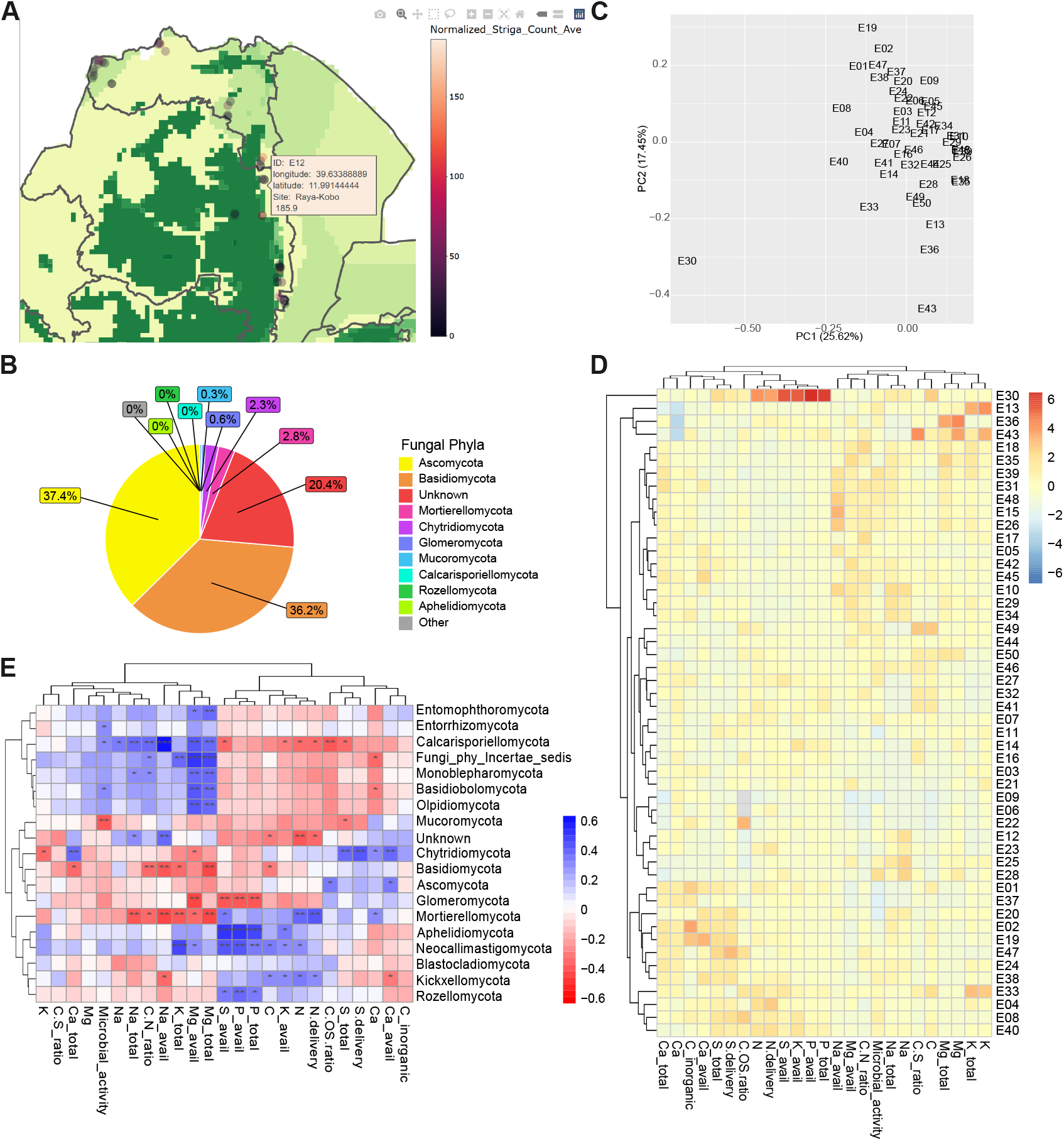
Demonstration of Interactive Application. Users can browse through the soil data using an interactive map **(A)** and get summaries of physio-chemical and microbial compositions for individual sample points **(B)**. Data visualization examples of Striga infestation level and Fungal phyla composition from soil sample E12 is represented in A & B. Users can explore the relationship between samples based on user-selected data in analyses like PCA **(C)** and generate heatmaps **(D)**. In addition to being able to reproduce figures from the paper, users can investigate other correlations between different microbial taxa levels and user-selected soil traits **(E)**. Plots were saved directly from the app then altered to fit figure panels here.

## Discussion

Numerous factors play a role in the distribution and devastating effects of Striga on crop production in sub-Saharan Africa. This study focused on a smaller, yet highly relevant region within Africa, representing multiple agro-climatic zones and Striga variability. Further, we quantified 32 physico- chemical parameters and microbial community composition, representing a diversity of soil composition from farmer’s and other fields under variable Striga seedbank and infestation levels in the sorghum belt of Ethiopia. Counts of emerged Striga plants were recorded during the soil collection, however, agricultural management strategies were not controlled and may artificially influence the appearance of Striga. As described previously, assessment of Striga presence through soil seedbank represents a measurement that may be more reflective of actual Striga occurrence in working fields than recording visible Striga plants and is a preferred Striga recording strategy [17]. Conversely, qPCR analysis of the seedbank is liable to overestimation due to non-viable or degraded Striga seeds still present in the soil. Regardless, our data shows a solid correlation between Striga seedbank and infestation (r = 0.67, p = 0.0001) which supports their comparable nature for assessing Striga occurrence [17]. The soil data also reflects high heterogeneity of Striga presence within fields, resulting in relatively high standard error measurements for both biological replicates of infestation and technical soil replicates of seedbank. Hence, accurate and extensive recording of Striga seedbanks and infestation in active agricultural regions proves to be a complex task that hinders monitoring of the effectiveness of existing and new mitigation practices.

Previous continental African and global-scale modeling studies have identified soil properties like nitrogen and clay content as important predictors of *Striga* habitat suitability [23]. These patterns are thought to be partly driven by the fact that host plants increase strigolactone secretion, promoting Striga germination, in nutrient-poor soils, particularly under phosphorus (P) or nitrogen (N) limitation typical of low-input farming systems [4,24]. Therefore, one might expect higher soil nutrient levels to reduce *Striga* infestation [24–27]. However, in our study, we did not observe significant correlations between soil P or N and *Striga* occurrence. Instead, we found that clay, potassium (K), sulfur (S), calcium (Ca), and carbon (C) were negatively correlated with infestation (**Figure 2B**). This divergence may reflect differences in nutrient ranges of study areas or nonlinear relationships with Striga. For example, Bellis et al. (2020) found *Striga* habitat suitability peaked at intermediate nitrogen levels (400–1000 mg/kg), while our soils ranged more widely (0–3970 mg/kg), potentially obscuring similar trends when fitted with linear models. Regional studies further support the complexity of these relationships. For example, in the Tigray region of Ethiopia, Gebreslasie et al. (2016) observed a negative correlation between *Striga* and available P, but no clear relationship with total N [25]. In Nigeria, Ekeleme et al. (2014) reported region-specific variation with Bauchi State showing *Striga* negatively correlated to clay, exchangeable K, and Ca, aligning with our results; while conflicting trends emerged in Kano State, underscoring geographic heterogeneity [26]. Dugje et al. (2008) also highlighted how relationships between nutrients and *Striga* vary by region in Nigeria [27]. Nutrient-based variation sustains the need to also consider soil texture, which plays a key role in modulating soil moisture, nutrient retention and microbiological communities [9,28] Clay content, specifically, has been repeatedly identified as an important variable in *Striga* ecology. Studies in Nigeria observed negative relationships between clay and *Striga* density [26,27], while in Ethiopia, positive associations with sand are reported, consistent with our finding of an negative relationship with clay [25]. Complexities in environmental context and interactions among soil properties may explain inconsistent findings across studies and highlights the need for further investigation into how soil dynamics affect Striga pressure.

While amplicon sequencing is typically not precise enough to capture the intraspecific diversity within a given genus, it was used to record unknown microbes as well as broader taxonomic trends. We identified a number of rare or taxonomically unknown taxa microbes where presence versus absence could be impactful to the Striga-sorghum interaction. Soils across the area studied demonstrated many of these rare ASVs which could have been better represented with a larger sample set. Bacterial diversity does not greatly vary across the sampled field soils, but fungal diversity did (**Figure 2B, Supp. Data 3**). This observation was supported by the sensitivity analysis which demonstrated that the fungal community is the most responsive to changes in Striga occurrence (**Figure 3A-B**). We next determined if specific fungal and bacterial taxa were correlated with Striga, and further, if any could be associated with lower Striga infestation. Several fungal and bacterial genera were found to either positively or negatively correlate with Striga infestation and seedbank. Of note is the phylum Glomeromycota which houses arbuscular mycorrhizal fungi, known for its beneficial interactions with plants under nutrient poor conditions. This phylum negatively correlated with levels of phosphorus in the soil, while genera within this taxa positively correlated with Striga (**Figure 4B, Figure 5E, Supp. Data 4**). Targeted analysis of a strain belonging to the genus *Neocosmospora* validated the hypothesis that a fungal isolate can increase Striga germination, indicative of the potential of this fungal genus to induce suicidal germination (**Figure 4D**). Efforts to understand the links between microbial communities and plant interactions will be continued and collaborated on to create large-scale data on the functional potential of soil microbes to interfere with the life-cycle of this and other devastating root parasitic weeds.

## Materials and Methods

### Soil Collection

A total of 50 soil samples were taken from naturally and artificially Striga infested fields in Amhara (Kemise, North Shewa, South and North Wollo Zones) and Tigray (West, Central and South zones) regions of Ethiopia in October, 2017 as described within Mitiku et al 2022. Forty-eight soil samples were collected from naturally infested agro-ecological zones as well as two from artificially infested fields (Figure 1). Sorghum fields with four categories (zero, low, medium and high) of Striga field infestation were randomly selected, and 4 samples were taken from the top layer (0-20 cm) around the root zone of a sorghum plant then combined to form one composite sample per field. Exceptions were samples E10, E15, E42, E44, E45, & E48, where sorghum plants were not uniformly distributed and only 1 sample was taken. Striga infestation and the Striga seedbank were quantified for forty-eight naturally infested soils (**Figure S1, Figure 2C-D**). Physico-chemical parameters were determined from forty-eight soils (**Figure 2A, S2A**).

### Measurement of Striga Infestation and Seedbank

Striga emergence was counted from four randomly chosen spots per field site, and the number of sorghum plants counted per m^2^ was used to normalize the number of emerged Striga plants. Striga emergence data were not obtained for samples E15 and E48 because these were from artificially Striga-infested fields. Seed bank density was assessed in Mitiku et al. 2022 using a qPCR approach.

### Soil Physical and Chemical Parameters

Soil physico-chemical analysis was done at Eurofins Agro (Eurofins Agro, Wageningen, The Netherlands) (**Supp. Data 1**) [18] The 4 macronutrients required in the highest amount by plants (N, K, Ca, Mg) were also described in percentages of nutrient make-up. H and Al were also measured for this purpose, but all samples were below the minimum detection threshold and discarded from further analysis. Datasets used in physico-chemical and Striga analyses excluded artificially infected fields (Samples E15 & E48) as well as an outlier sample, E30.

### Microbial Community Analysis

The composite bacterial and fungal microbiome were quantified for forty-six soils, excluding two soils which had been planted a separate experiment involving intercropping systems (E49 and E50) (**Figure S2B-C**). Bacterial DNA preparation, sequencing, and processing are explained in Abera et al. (2022) [18]. The same DNA samples were used for fungal sequencing targeting the ITS2 region of Internal Transcribed Spacer (ITS) gene, using primers 58A1F (5′- GCATCG ATGAAGAACGC-3′) and ITS4 (5′- TCCTCCGCTTATTGATATGC-3′). Sequencing was performed at BaseClear B.V. (Leiden, Netherlands).

Fungal isolates from Lombard et al. (2025) and Macia-Vicente et al. were collected from the same soils analyzed in this study. Fungal ASV sequences were aligned with the sequences from these soils deposited in GenBank [accession numbers PX349626 - PX351653] using nucleotide BLAST+ [21,29– 31]. Matches were filtered to the highest bit score. The ASVs were then reassigned the genera of their best match isolate. In the event an ASV sequence matched multiple accessions, the ASV was assigned the genera that represented at least 70% of its matches (**Supp. Data 5**). Microbial data followed summation at different taxa levels, zero imputation with a constant value and centered log-ratio transformation from package Tjazi (0.1.0.0) [32] using the “logunif” method at 1000 repetitions. ASV data was rarefied and alpha diversity measures were calculated using phyloseq (1.46.0).

### Generalized Joint Attribute Modeling (GJAM)

To integrate microbiome data with Striga infection and soil factors, we used generalized joint attribute modelling (GJAM) via gjam package version 2.6.2 in R v4.3.1 [19]. To evaluate the role of Striga infection and Striga seedbank on the microbial communities (bacteria and fungi) we grouped those variables into discrete categories to represent Zero, Low, Medium, and High levels of both Striga infestation and Striga seedbank. Then two models were generated to evaluate (1) the effects of Striga Infestation and (2) the effects of Striga Seedbank (**Supp. Data 3**). GJAM is a Bayesian model that estimates the regression coefficient via Gibbs sampling. In our case, we used 20,000 iterations before reaching stability of the regression coefficients. Composition count was selected as data type since the microbiome data is to be considered compositional based on heterogeneous number of reads per sample (Gloor et al. 2017). The impact of the categories of Striga Infestation (model 1) and Striga Seedbank (model 2) in shaping the microbiome and soil factors was evaluated per groups of variables via Sensitivity analysis as in Rotoni et al., (2022) [33]. The sensitivity to predictors analysis evaluate how the different explanatory variables (in our case the different levels of striga seedbank and infestation) influenced the different groups of variables, thus allowing us to identify the levels of striga seedbank and infestation that resulted in the biggest shifts in the microbial community and soil factors.

### Correlation Analyses

Striga data followed imputation with a constant value of 1 then log 10 transformations to be used in Pearson correlations against physico-chemical and microbial soil data using the R package “psych” (2.4.3). Soil samples missing measurement categories (i.e. E15 and E48 which did not have Striga measurements, E49 and E50 which did not have microbial data) were excluded from analyses requiring that data. E30 was excluded from all physico-chemical related analyses for being an outlier. Bonferroni p-value adjustment was applied to results (**Supp. Data 3-4**).

### *In vitro* Striga Assays

Sterilized Striga seeds collected from Qilee in 2022 were imbibed in 1% water agar and incubated 6 days at 30C in the dark. Seeds were treated with the following: no fungus & no GR24+, fungus & no GR24, no fungus & GR24, fungus & GR24. Fungi were inoculated by applying a suspension of 1000 conidia/replicate, after which plates were incubated at 30C in the dark for 6 days. Seed were then treated/untreated with 1 mg/ml GR24, and after 24 h the percentage of seed germination per treatment scored (**Supp. Data 5**). Treatments consisted of 24 replicates, each with 20–80 seeds.

### Interactive Application

The web application was made using the shiny package (1.8.0) in R. It contains all the soil data included in this article (physico-chemical soil analysis, agro-climatic zones, summary Striga measurements, and ASV sequencing results) (**Supp. Data 1-2**). The app is equipped with similar analyses as presented in this study with additional ways of visualizing the data. Maps in the paper and app were created using packages ggplot2 (3.5.1), plotly (4.10.4) and sf (1.0-15) and polygons from the github repository of user georgique and https://gadm.org/. Correlation analyses follow R package “psych” (2.4.3), PCA analysis uses the base stats (4.3.1) package, and scatter plots are produced using base stats and graphics (4.3.1). Random forest and decision tree analysis (randomForest 4.7-1.2) is included for demonstrative purposes of future use, using all selected samples in the training dataset and is not equipped for a testing dataset to assess accuracy. Scripts for this ShinyApp can be found in the following github repository: https://github.com/taffymieh/Soil-App.

## Supporting information

Supplemental Data 1: Soil geographical, agro-climatic, physico-chemical and summarized microbial data.

Supplemental Data 2: Microbe sequencing data.

Supplemental Data 3: Correlation analyses.

Supplemental Data 4: Taxon level correlation analysis.

Supplemental Data 5: Reassigned fungal genera and in vitro assay.

## Acknowledgements

This work was supported, in whole or in part, by the Gates Foundation grant to Netherlands Institute of Ecology (NIOO) “Engineering Soil and Plant Microbiomes for Enhanced Crop Productivity in Africa (OPP1082853) and “PROMISE II: Promoting microbes for integrated Striga eradication” (INV-047255). The conclusions and opinions expressed in this work are those of the author(s) alone and shall not be attributed to the Foundation. Under the grant conditions of the Foundation, a Creative Commons Attribution 4.0 License has already been assigned to the Author Accepted Manuscript version that might arise from this submission. Please note works submitted as a preprint have not undergone a peer review process.

## Contributions

TTa, GB, DE, TM, EMP, JGMV, TTe, PC, EEK, JMR, SMB contributed to experimental design. GB, DWE, LL, LAG, DR, TM, EMP, JGM-V, DYL, UT, JD, RD, RRL, TTe contributed to data acquisition. TTa, GB, MFAL, LL, LA-G, SS, TM, EMP, JGM-V, RD, EEK, PWC, JMR, SMB contributed to data analysis. TTa, BG, MFAL, TM, EMP, JGM-V, EEK, JMR, SMB contributed to writing.

## SUPPLEMENTAL DATA

**Supplemental Data 1**: Soil geographical, agro-climatic, physico-chemical and summarized microbial data.

1. Physico-chemical: Soil identifier (ID); location (site); Altitude (meters); latitude (degrees); longitude (degrees); pH; soil sample chemical analysis of total (_total) and available (_avail) elements, N (nitrogen), S (sulfur), P (phosphorus), K (potassium), Ca (calcium), Mg (magnesium); carbon (C, %); soil sample physical analysis (%) of organic matter (organic), inorganic carbon (C_inorganic), carbonated lime (carbonated_lime), clay (<2 um), silt (2-50 um), and sand (>50 um); soil nutrient profiling (%) of calcium (Ca), magnesium (Mg), potassium (K), sodium (Na); nitrogen delivery (N.delivery); sulfur delivery (S.delivery); carbon:nitrogen ration (C/N_ratio); carbon:sulfur ratio (C/S_ratio); hydrocarbon concentration between original and tested soil (C/OS_ratio); electrical exchange measurements of Cation Exchange Capacity (Clay_humus, mmol+/kg); microbial activity (mg N/kg); and Koeppen-Geiger climate type (Agro_Climate_type).
2. Striga Measurements: Pooled Striga measurements of seedbank and field infestation and their log (base 10) transformations used in analysis.
3. Field Infestation: original Striga field infestation measurements normalized by sorghum presence.
4. Seedbank: Original Striga seedbank measurements by qPCR (Striga seeds per 150g).
5. PCA Eigens Physico-chemical: PCA eigen values for variables in physico-chemical analysis obtained by extracting the loadings from the PCA result of prcomp(). PCs 1-30 to explain 100% of the variation.

**Supplemental Data 2**: Microbe sequencing data.

1. 16S Sequencing ASV counts: Bacterial and Archea ASV counts by 16S sequencing with technical replicates (samples E5, E33, E36, and E48) averaged.
2. ITS Sequencing ASV Counts: Fungal ASV counts by Internal Transcribed Spacer sequencing with technical replicates (samples E5, E33, E36, and E48) averaged.
3. Bacteria Diversity: Diversity indexes calculated on rarefied ASVs by the phyloseq package.
4. Fungus Diversity: Diversity indexes calculated on rarefied ASVs by the phyloseq package.
5. PCA summary: Importance of components from Bacterial and Fungal PCA analysis.
6. Bacteria Loadings: PCA loadings for ASVs and their taxonomy.
7. Fungal Loadings: PCA loadings for ASVs and their taxonomy.

**Supplemental Data 3:** Correlation analyses.

1. GJAM Infestation: Sensitivity values from GJAM analysis for soil measurements grouped as bacteria, fungus, soil factors, and Striga calculated for ranges of Striga infestation.
2. GJAM Seedbank: Sensitivity values from GJAM analysis for soil measurements grouped as bacteria, fungus, soil factors, and Striga calculated for ranges of Striga seedbank.
3. Summary Correlation R values: R value results of all pairwise Pearson correlation analysis on physico-chemical, microbe diversity and log (base 10) transformations of Striga seedbank (average and SE) and infestation (average and SE).
4. Summary Correlation P values: Significance of all pairwise Pearson correlation analysis on physico-chemical, microbe diversity and log (base 10) transformations of Striga seedbank (average and SE) and infestation (average and SE). P value in bottom triangle, Bonferroni adjusted p values in top triangle.

**Supplemental Data 4:** Taxon level correlation analysis.

ASV data was aggregated at each taxon level, imputed and CLR transformed. Unidentified taxa were grouped into an “Unknown”. Pearson correlation was run with each dataset against log (base 10) transformations of Striga seedbank (average and SE) and infestation (average and SE). Structure of each sheet is the same. Contains R value, P value, and Bonferroni adjusted P value.

1. Bacterial Phyla
2. Bacterial Class
3. Bacterial Order
4. Bacteria Family
5. Bacterial Genera
6. Fungal Phyla
7. Fungal Class
8. Fungal Order
9. Fungal Family
10. Fungal Genera

**Supplemental Data 5:** Reassigned fungal genera and in vitro assay

1. ASV Accession Matches: Best sequence match results between ASV sequences and GenBank accession records of fungal cultures isolated from these same soils.
2. New Fungal Genera Correlation: Pearson correlation analysis results on Fungus genera based on taxonomy from matching ITS sequence in fungal strain collection and log (base 10) transformations of Striga seedbank (average and SE) and infestation (average and SE). Contains R value, P value, and Bonferroni adjusted P value.
3. Assay Data: Treatments and Striga germination measurements from testing fungal strain interaction with striga seeds.

