## Supplemental Figures 1-3 for "Disentangling the importance of microbiological and physico-chemical properties of Ethiopian field soils for the Striga seed bank and sorghum infestations"

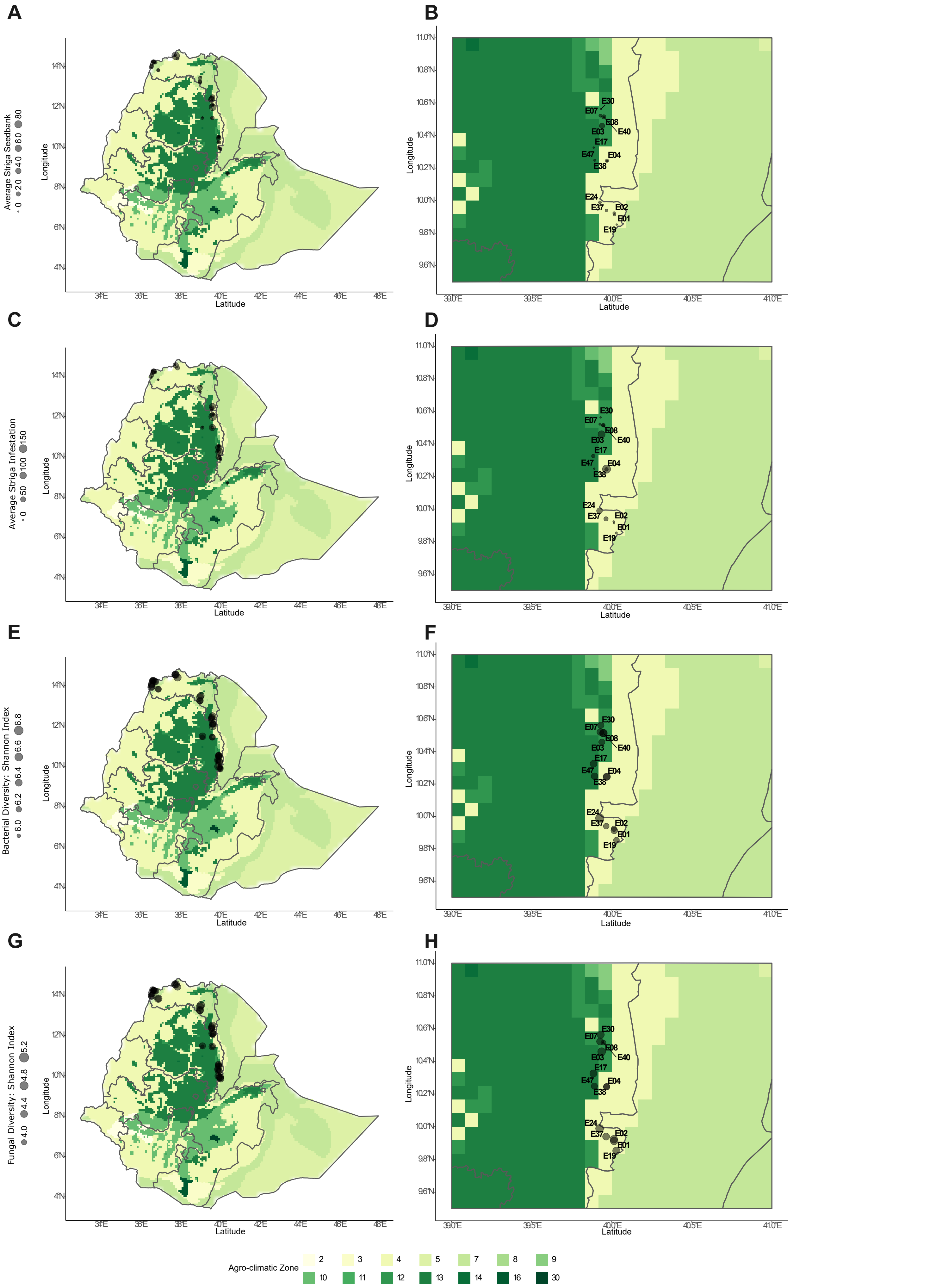


**S Fig1. Striga occurrence and microbiological diversity vary across soil samples.**
Striga seedbank and field infestation levels varied across the sampling sites **(A & C)**, even between sites of close proximity such as in the Kewet area **(B & D)**. The shannon index, a metric of alpha diversity was calculated separately for fungal and bacterial microbes and also varied between sites **(E-H).**

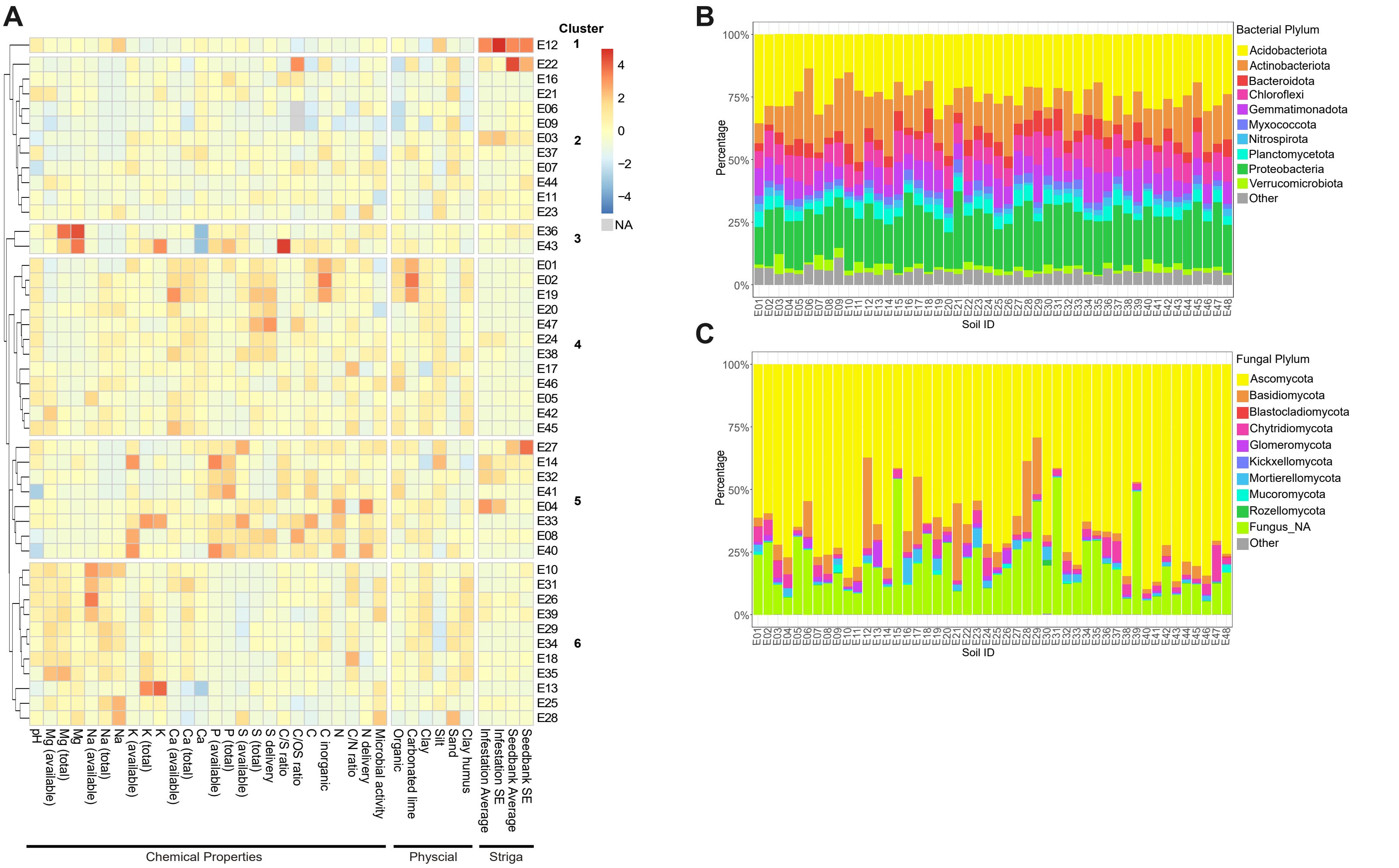


**S Fig2. Some soil factors negatively correlate with Striga seedbank or field infestation.**
**A.** Heat map of soil physico-chemical properties and Striga measurements of all naturally infested soils excluding outlier sample 30. Color represents the relative value across samples (pheatmap). Soil samples were sorted by hierarchical clustering using a Euclidean distance metric with a tree cut off at 6 branches. **B.** Proportions of the most abundant bacterial phyla within each soil sample. **C.** Proportions of the most abundant fungal phyla within each soil sample.

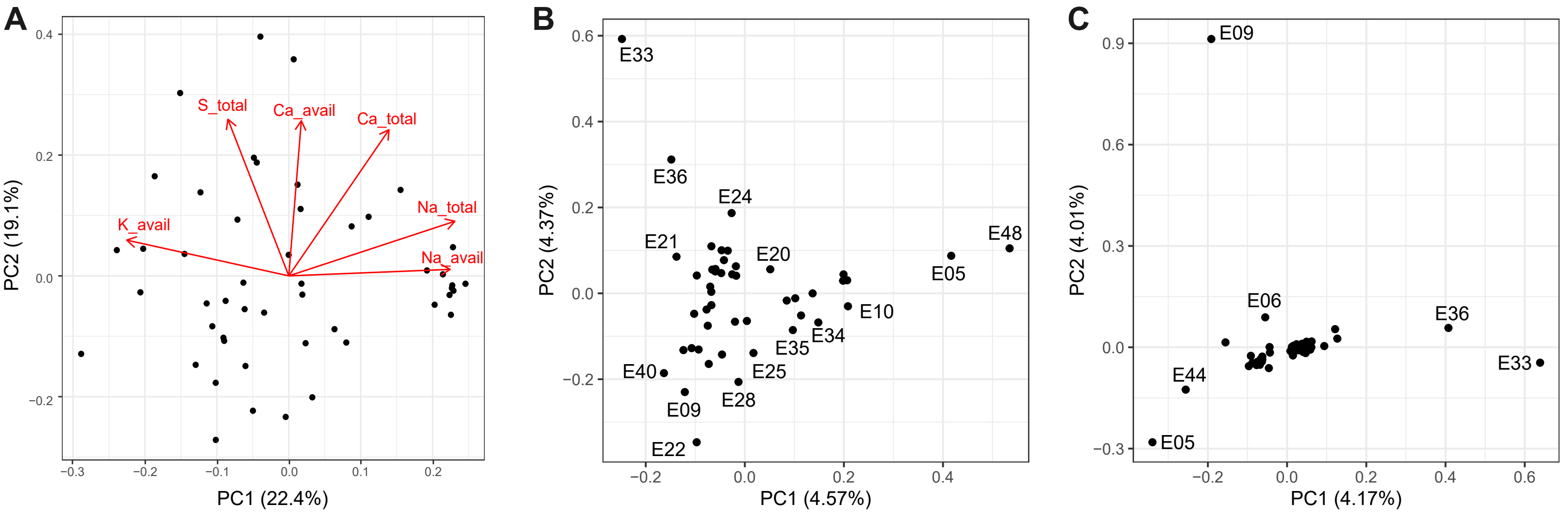


**S Fig3. Soil sample variability is driven by many factors.**
PCA (Principle Component Analysis) of physico-chemical composition (**A**), bacterial ASV community (**B**), and fungal ASV community (**C**) of Ethiopian soil samples. Red arrows represent loading vectors of the 3 most informative factors driving variation in each PC1 (Na_total, K_avail, Na_avail) and PC2 (S_total, Ca_avail, Ca_total) for the physico-chemical composition.
